# Isolation and preliminary characterization of extracellular vesicles from bottlenose dolphin (Tursiops truncatus) and long-finned pilot whale (Globicephala melas) blow

**DOI:** 10.64898/2026.05.29.728645

**Authors:** Valentina Moccia, Cinzia Centelleghe, Andrea Zendrini, Selene Tassoni, Luca Ceolotto, Bertrand Bouchard, Eva Alvarez, Gaia Pesce, Paolo Bergese, Annalisa Radeghieri, Sandro Mazzariol, Valentina Zappulli

## Abstract

Cetaceans are key sentinel species for environmental health monitoring. Although sampling from free-ranging animals is challenging, the analysis of cetacean blow offers a minimally invasive approach to assess their health status. Extracellular vesicles (EVs) are cell-derived nanostructures present in biological fluids and widely studied as disease biomarkers in humans. Despite the potential for similar uses, EVs have not been studied in cetacean blow to date . This proof-of-concept study aims to assess the feasibility of the isolation and characterization of EVs from blow samples collected from five bottlenose dolphins (*Tursiops truncatus*) kept under human care and from a free ranging specimen of long-finned pilot whale (*Globicephala melas*). EVs were purified from bottlenose dolphin blows by ultracentrifugation (UC) or size exclusion chromatography (SEC) and from the long-finned pilot whale by SEC. Particle concentration and size distribution were assessed by Nanoparticle Tracking Analysis (NTA), morphology by Air-atomic force microscopy (AFM) and protein expression by Western Blotting (WB). NTA revealed a higher mean particle concentration in bottlenose dolphin EVs isolated by UC compared to SEC, while EVs isolated from the long-finned pilot whale presented a lower particle concentration. AFM confirmed the presence of EV-like particles within the typical EV size range in bottlenose dolphin’s EVs obtained both by SEC or UC. All EV samples were positive for CD9 and integrin-β and negative to Calnexin. SEC was more sensitive to detect OmpA, a membrane protein of Gram-negative bacteria, in EVs from both species. Our pilot study demonstrates that EVs are present in cetacean blow and can be isolated and characterized. Future investigations focused on characterizing and quantifying a wider array of EV associated molecules may further the application of blow EV analysis for cetacean health assessments.

## Introduction

Cetaceans are recognized as good species for monitoring marine ecosystem health due to their long lifespans, high lipid content and position at the top of the trophic level [1–8] which lead to bioaccumulation and biomagnification of xenobiotics [2,5–8]. In addition, as mammals that share comparable food sources with humans (e.g., fish and cephalopods), the evaluation of cetacean conditions can also provide information useful for public health [9]. Despite their relevance, assessing the health of free-ranging cetaceans remains arduous. Usually, most investigations tend to rely on *post-mortem* analyses, which limit the evaluation of their health conditions *in vivo*. More broadly, evaluating the health status of wild animals presents conceptual and practical challenges. In animals, health has been described as “a state of physical and psychological well-being and of productivity, including reproduction,” and is typically inferred from measurable parameters such as food intake, fecal output, and body condition [10–12]. A more contemporary perspective further defines health as “the ability to adapt and to self-manage,” which is the capability of maintaining physiological homeostasis under changing conditions [13]. Within this framework, disease can be defined as a disruption of one or more physiological functions, which can also arise from interactions with infectious or non-infectious pathogens. Importantly, pathogenicity is not an intrinsic property of a microorganism alone but depends on the dynamic interplay between host susceptibility and microbial characteristics [12]. For wild species, where baseline data and pathogen information are often limited, the development of novel, minimally invasive tools for health assessment is therefore particularly critical. In this context, in the last two decades, the sampling and analysis of cetacean blow have emerged as an alternative for the study of their biological conditions [14–18]. In contrast to tissue or blood sampling, blow can be collected with a minimally-invasive technique from free-ranging individuals using Unmanned Aerial Vehicles (UAVs, or drones) [16,17]. Cetacean blow contains a mixture of surfactant, mucus, cells and cellular debris originating from the lower respiratory tract [17,19,20]. This provides useful information about the respiratory microbiome [15,16,21–23], hormonal concentrations [14,24,25] and the immune status across different species of cetaceans [20,26,27]. However, a detailed analysis of other components with diagnostic potential has not yet been conducted for this biofluid.

It has recently been demonstrated that beyond their cellular and soluble molecular components, biofluids (e.g. blood, urine, cerebrospinal fluid, saliva, exhaled breath condensate) contain a diverse population of nanoparticles [28]. Among these, extracellular vesicles (EVs) are particularly relevant as potential health biomarkers [29–31]. Extracellular vesicles are membrane-bound structures released by all cells into the extracellular space [28,32] carrying different bioactive compounds [28,33]. The composition of EVs differs according to the cell type where they were synthesized and to their physiological – or pathological – state [34]. After their release, EVs allow intra- and inter-organismal communication, acting as mediators of cellular communication and regulators of various bioprocesses [35–39]. Because of their availability in every biofluid and of their complex composition, EV-analysis is now commonly used to detect novel biomarkers of diverse human diseases and potential indicators of exposure to xenobiotics in humans [40–42]. The potential of EVs for disease diagnosis started to be assessed for veterinary medicine [29,43–46]. However, probably due to the difficulties of sampling, comparable efforts are still scarce for wildlife [47]. For cetaceans, only preliminary isolation and characterization of EVs released from cultured fibroblasts of bottlenose dolphins (*Tursiops truncatus*) and Cuvier beaked whales (*Ziphius cavirostris*) [48] and serum from five species (minke whale *Balaenoptera acutorostrata*, fin whale *B. physalus*, humpback whale *Megaptera novaeangliae*, orca *Orcinus orca*, and Cuvier’s beaked whale) have been conducted [49].

Given that EVs have been identified in every biological fluid examined so far, and considering the potential of blow analyses for cetacean health assessment, we aimed to isolate and characterize EVs in cetacean blow. We used two different EV purification methods – one based on ultracentrifugation (UC) and one on size exclusion chromatography (SEC) on blow samples collected from five captive bottlenose dolphins. Our methods were further validated on a blow sample collected from a free-ranging long-finned pilot whale (*Globicephala melas*). This is the first demonstration that EVs can be isolated from blow samples.

## Materials and methods

### Blow sampling

Blow samples were collected from five bottlenose dolphins (*Tursiops truncatus,* Table 1) maintained under human care at aquarium of Genova (Costa Edutainment S.p.A., Genova, Italy). An additional blow sample was collected from one free-ranging long-finned pilot whale (*Globicephala melas*) during a fieldwork expedition three nautical miles from the French coast (Mediterranean sea, 43°02’13.4“N 5°51’53.2”E). The collected samples were transported and stored at the Mediterranean Marine Mammals Tissue Bank of the Department of Comparative Biomedicine and Food Science (University of Padua, CITES authorization IT020).

**Table 1.**
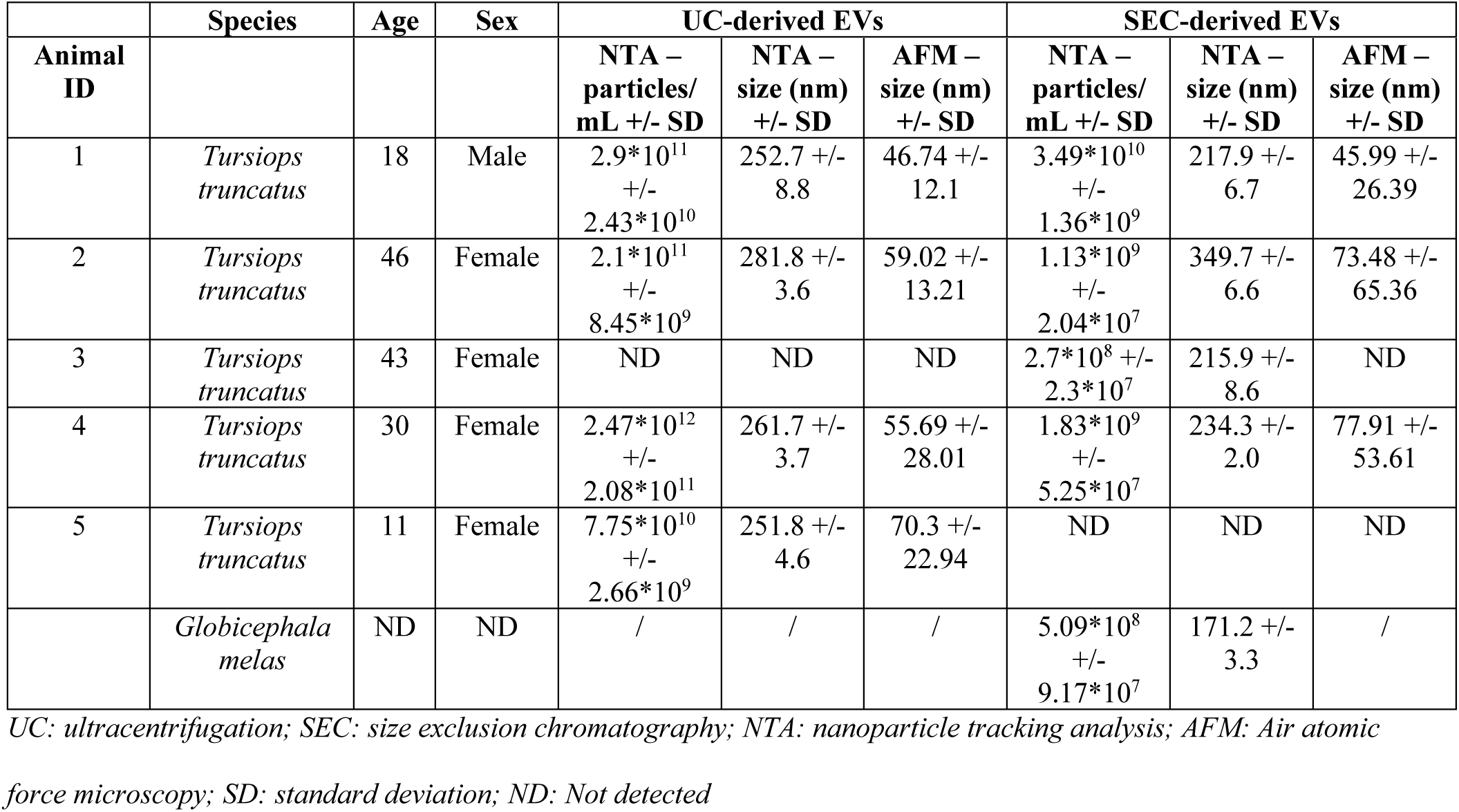
ID number, age, sex and characterization results of extracellular vesicles isolated from the blow of five bottlenose dolphins (*Tursiops truncatus*) and from long-finned pilot whale (*Globicephala melas*)

|  | Species | Age | Sex | UC-derived EVs | SEC-derived EVs |
| --- | --- | --- | --- | --- | --- |

| Animal ID |  |  |  | NTA – particles/mL +/- SD | NTA – size (nm) +/- SD | AFM – size (nm) +/- SD | NTA – particles/mL +/- SD | NTA – size (nm) +/- SD | AFM – size (nm) +/- SD |
| --- | --- | --- | --- | --- | --- | --- | --- | --- | --- |
| 1 | *Tursiops truncatus* | 18 | Male | 2.9*10<sup>11</sup> +/- 2.43*10<sup>10</sup> | 252.7 +/- 8.8 | 46.74 +/- 12.1 | 3.49*10<sup>10</sup> +/- 1.36*10<sup>9</sup> | 217.9 +/- 6.7 | 45.99 +/- 26.39 |
| 2 | *Tursiops truncatus* | 46 | Female | 2.1*10<sup>11</sup> +/- 8.45*10<sup>9</sup> | 281.8 +/- 3.6 | 59.02 +/- 13.21 | 1.13*10<sup>9</sup> +/- 2.04*10<sup>7</sup> | 349.7 +/- 6.6 | 73.48 +/- 65.36 |
| 3 | *Tursiops truncatus* | 43 | Female | ND | ND | ND | 2.7*10<sup>8</sup> +/- 2.3*10<sup>7</sup> | 215.9 +/- 8.6 | ND |
| 4 | *Tursiops truncatus* | 30 | Female | 2.47*10<sup>12</sup> +/- 2.08*10<sup>11</sup> | 261.7 +/- 3.7 | 55.69 +/- 28.01 | 1.83*10<sup>9</sup> +/- 5.25*10<sup>7</sup> | 234.3 +/- 2.0 | 77.91 +/- 53.61 |
| 5 | *Tursiops truncatus* | 11 | Female | 7.75*10<sup>10</sup> +/- 2.66*10<sup>9</sup> | 251.8 +/- 4.6 | 70.3 +/- 22.94 | ND | ND | ND |
|  | *Globicephala melas* | ND | ND | / | / | / | 5.09*10<sup>8</sup> +/- 9.17*10<sup>7</sup> | 171.2 +/- 3.3 | / |

UC: ultracentrifugation; SEC: size exclusion chromatography; NTA: nanoparticle tracking analysis; AFM: Air atomic force microscopy; SD: standard deviation; ND: Not detected

For bottlenose dolphin sampling, blow samples were collected from each dolphin in duplicate. Briefly, blow samples were collected using a six-well plate positioned 40 cm above the blowhole immediately after exhalation in respond to trainer’s command. Blow samples were dissolved in 1 mL of 0.2 µm double-filtered PBS (dfPBS) and transported and stored at + 4°C until EV-isolation, which was performed within 24 hours from collection.

The long-finned pilot whale was sampled using a six-well plate that was secured face down using a custom-built holder attached to a drone (SwellPro SplashDrone 4) (S1 Video). The drone was flown by a licensed biologist and positioned at approximately three meters above the whale (following several attempts to determine the best distance; data not shown). This allowed exhaled breath condensate droplets to adhere to the plate [17]. Immediately after collection, the wells were sealed to prevent contamination and evaporation. At the end of the sampling, samples were dissolved in 1 ml of dfPBS and stored at + 4°C during transportation to the laboratory (72 hours) where it was stored at -80 °C until EV-isolation. Environmental samples were collected during sampling of bottlenose dolphins. These were obtained by placing a six-well plate 70 cm above the surface of the animals’ pool to assess potential air and/or water contamination. Environmental samples were conserved and processed exactly as done for the blow samples.

### Extracellular vesicle isolation

Prior to EV purification, each 1 mL dfPBS-dissolved blow sample was centrifuged at 300×*g* then at 2000×*g* for 10 min at 4 °C and the pellet discarded after each step to remove any cell/cell debris. Then, blow-derived EVs were isolated with two different protocols, one based on UC and one based on SEC. Due to the small sample volume of the long-finned pilot whale’s blow, EVs were isolated only by SEC . For EV isolation by UC, the supernatant resulting from the 2000×*g* centrifuge was diluted in dfPBS to a final volume of 5 ml, transferred to ultracentrifuge tubes (Beckman Coulter, Brea, CA, USA) and ultracentrifuged at 100,000×*g* for 90 min at 4 °C (Optima L-90 K, Beckman Coulter) in a swinging bucket rotor (SW55ti, Beckman Coulter). The supernatant was discarded, and the EV-enriched pellet was resuspended in 100 μL of dfPBS for characterization analysis.

To perform EV-isolation with SEC, the supernatant resulting from the 2000×*g* centrifuge was concentrated with 100 kDa ultrafiltration tubes (Amicon Ultra centrifugal filters, Merck Millipore, Burlington, MA, USA) and loaded on the top of qEVoriginal/70 nm columns (IZON Science, Christchurch, New Zealand). SEC was then performed according to manufacturer’s instructions and fractions #7, #8, #9, and #10, containing most EVs with least co-isolated contaminants based on manufacturer’s guidelines were collected, pooled and concentrated with a 100 kDa ultrafiltration tube (Amicon Ultra centrifugal filters, Merck Millipore). Concentrated pooled fractions were finally collected and resuspended in 100 μL of dfPBS to perform further analysis.

### Nanoparticle Tracking Analysis

After EV isolation, EVs obtained with UC or SEC from each blow sample were evaluated for particle concentration and size distribution by Nanoparticle Tracking Analysis (NTA) with NanoSight NS300 (Malvern). Samples were progressively diluted in dfPBS to reach the correct dilution to gain reliable measurements. For each sample, camera level was set at 12, and three movies of 60 s each were recorded and analyzed using the 3.4 NTA software. For particle quantification, the detection threshold was set at 5 and measurements were considered reliable when within the following instrument optimal working ranges: particles per frame from 20 to 120; particle concentration between 10^6^ and 10^9^ per mL; ratio of valid particles to total particles higher or equal to 1:5.

To test the differences between groups (UC and SEC), a statistical analysis was performed with Mann-Whitney test using GraphPad Prism 9 software. The level of significance was fixed as *p* < 0.05.

### Atomic Force Microscopy

The solvent in EV samples that had been obtained from the bottlenose dolphin blow samples was exchanged for Milli-Q water (Merck Millipore Ltd., Darmstadt, Germany) using Amicon Ultra-0.5 mL centrifugal filter units with a 30 kDa molecular weight cutoff (Merck Millipore Ltd., Darmstadt, Germany). The solvent exchange step was performed three times, centrifuging the samples at 12,000×g for 15 minutes each.

Following the final wash, EVs were resuspended in 100 μL of Milli-Q water and subsequently diluted 1:100 in the same solvent. A 5 μL aliquot of the diluted EV suspension was deposited onto freshly cleaved mica sheets (Grade V-1, thickness 0.15 mm, dimensions 15 × 15 mm²; Ted Pella Inc., Redding, CA, USA), and the samples were allowed to dry for 10 minutes at room temperature.

Air-atomic force microscopy (AFM) imaging was performed using a Nanosurf NaioAFM (Nanosurf AG, Liestal, Switzerland) operated in tapping mode and equipped with a Multi75-AI-G probe (Budget Sensors, Sofia, Bulgaria). Images were acquired with 1-0.6 seconds per line (P-Gain 5000, I-Gain 500, D-Gain 0). Image processing and analysis were carried out using Gwyddion software (version 2.67).

### Western Blot Analysis

To evaluate the presence of EV markers, Western Blotting (WB) was performed on each UC and SEC-derived EV sample. Proteins extracted from a bottlenose dolphin fibroblast cell line (Sea Sentinels System patent No. 102020000003248) were included as positive controls for EV and mammalian cellular markers, while an *Escherichia coli* lysate was included as a positive control for bacterial proteins. Prior to WB analysis, protein concentrations were calculated for each sample using a Pierce BCA protein Assay Kit (Thermo Fisher Scientific), according to the manufacturer’s protocol. Namely, 20 μg of proteins were denatured at 70 °C for 10 min and then resolved using NuPAGE 4–12% Bis–Tris gel (Thermo Fisher Scientific) before transferring to a nitrocellulose membrane. To block nonspecific binding sites, blots were incubated for 90 min in 7% non-fat dry milk in TBS-T (TBS containing 0.1% Tween-20) at room temperature. Then, blots were incubated at 4 °C overnight with rabbit or mouse primary antibodies against human Integrin-β1 (1:5000; GeneTex GTX128839, Irvine, CA, USA) and CD9 (1:500; Bio-Rad MCA694GT, Hercules, CA, USA), two membrane proteins, and calnexin (1:1000; Cell Signaling #2679, Danvers, MA, USA), a marker of the endoplasmic reticulum commonly used as negative control for EVs by demonstrating the absence of cell debris in the EV-preparation [50]. In addition, for further investigations on host-microbiome interactions, we also investigated the presence of bacteria or of their derived-EVs by incubating blots with a primary antibody against bacterial OmpA (1:1000; Biorbyt orb422682, Durham, NC, USA). All primary antibodies were diluted in TBS-T containing 3% non-fat dry milk. Then, membranes were incubated with a peroxidase-conjugated secondary antibody (1:3000; according to the species of the primary antibody, anti-Rabbit #32260 or anti-Mouse #32230, Thermo Fisher Scientific) diluted in TBS-T for 1 h at room temperature. Reactive bands were visualized using the SuperSignal West Pico PLUS Chemiluminescent Substrate detection kit (Thermo Fisher Scientific) with the iBright instrument (Thermo Fisher Scientific).

### Permits and authorizations

Blow sampling from bottlenose dolphins kept under human care was performed according to the Council Directive 1999/22/EC. All the samples were collected non-invasively during routine veterinary examinations, performed according to the Italian Decree 73/2005 and annexes, establishing the management objectives and prescriptions to maintain this species under human care in Italy. Blow sampling of the long-finned pilot whale was conducted under an official derogation authorizing the intentional disturbance of protected species (Arrêté préfectoral No. 84, Préfet maritime de Toulon, 04/07/2025).

## Results

### Nanoparticle size distribution, concentration and morphology

Both protocols (UC and SEC) allowed successful isolation of EVs from the five bottlenose dolphins, and the protocol (SEC) used on the long-finned pilot whale blow sample was also successful. However, NTA found that the EVs isolated from two bottlenose dolphin blow samples (UC-derived EVs of dolphin ID3 and SEC-derived EVs of dolphin ID5) and all of the environmental control samples were below the minimal detection limit, meaning that particle concentration was insufficient for AFM and WB.

For the remaining samples, particle concentration at NTA was heterogeneous among all dolphins’ blow samples, varying from 7.75*10^10^ (ID5) to 2.47*10^12^ (ID4) particles/mL in UC derived-EVs and from 2.7*10^8^ (ID3) to 3.5*10^10^ (ID1) particles/mL in SEC-derived EVs (Table 1). Despite the heterogeneity, the UC method yielded a higher mean EV particle concentration than the SEC method (Figure 1, *p* < 0.05, U = 0, n_1_ = 4, n_2_ = 4; 7.6*10^11^ +/- 1.1*10^12^ and 9.5*10^9^ +/- 1.7*10^10^ particles/mL, respectively). The measured particle size was similar between UC and SEC-derived EVs, with a mean size ranging from 251.8 +/- 4.6 nm (ID5) to 281.8 +/- 3.6 nm (ID2) for UC, and from 215.9 +/- 8.6 nm (ID3) to 349.7 +/- 6.6 nm (ID2) for SEC-derived EVs (Table 1).

**Fig 1.**
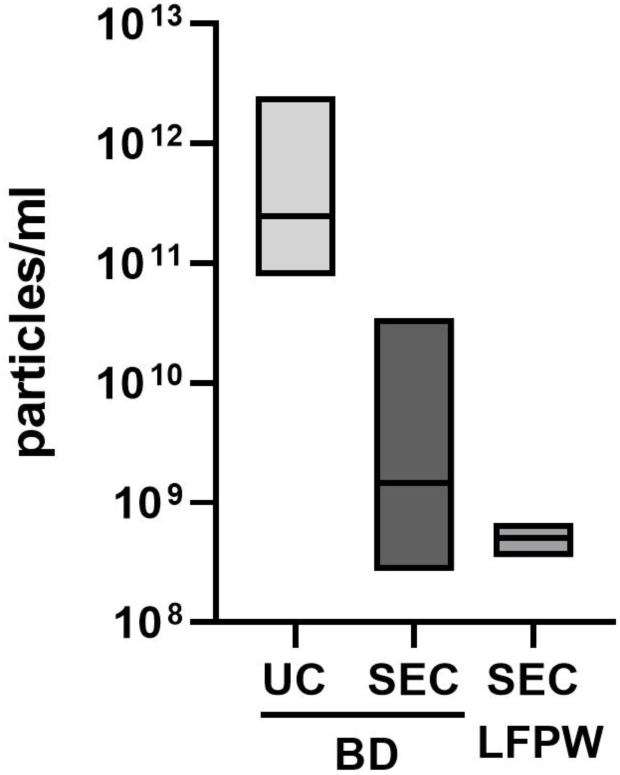
Comparison of the mean concentration of extracellular vesicles (EVs) isolated by ultracentrifugation (UC) or size exclusion chromatography (SEC) from bottlenose dolphin’s (BD) and long-finned pilot whale’s (LFPW) blow at Nanoparticle Tracking Analysis. The mean nanoparticle concentration was higher in BD UC-derived EVs (7.6*10^11^ +/- 1.1*10^12^ particles/mL) compared to BD SEC EVs (9.5*10^9^ +/- 1.7*10^10^ particles/mL, p<0.05) and to LFPW SEC EVs (5.09*10^8^ +/- 9.17*10^7^ particles/mL).

NTA also showed a particle concentration of 5.09*10^8^ +/- 9.17*10^7^ particles/mL with a mean size of particles of 171.2 +/- 3.3 nm of the EV sample isolated from the long-finned pilot whale blow with SEC (Table 1, Fig 1).

AFM was performed on EVs isolated from bottlenose dolphins’ blow to assess EV-presence through the examination of particles with a morphology resembling that of soft lipid particles and to verify the absence of large protein aggregates, which can frequently co-isolate with EVs [51]. The AFM analysis revealed round nano-objects within the typical dried-out EV size range [51] for both UC and SEC isolation protocols, with a mean diameter not statistically different (Fig 2a-c). Mean diameters of EVs for each isolation protocol are shown in Table 1. For sample ID3, which presented the lowest particle concentration at NTA, the number of particles detected was insufficient to perform the analysis. Protein aggregates and debris resulting from EV rupture were virtually absent in all of the samples (Figs 2b and 2c).

**Fig 2.**
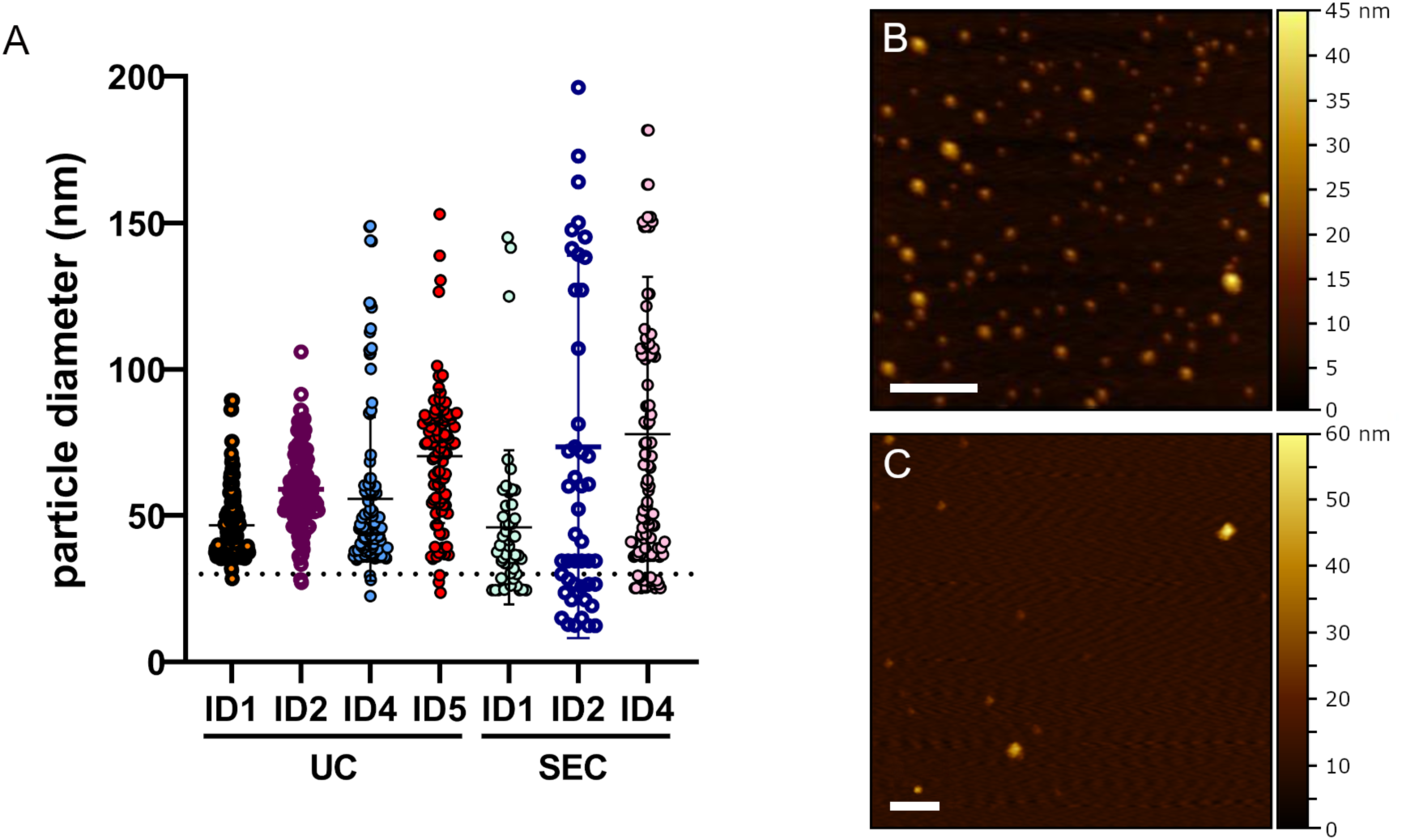
Results of Air-Atomic Force Microscopy (AFM) analysis performed on dolphin blow derived extracellular vesicles (EVs). Scatter plot showing the size of each particle measured at AFM for each EV sample isolated by ultracentrifugation (UC) or size exclusion chromatography (SEC). The dotted line represents the lower theoretically limit for EV-size (30 nm) (A); Morphology and size of EV-like particles isolated by UC from dolphin ID1 (B) and by SEC from dolphin ID1 (C).

### Protein expression

EV markers CD9 and integrin-β were detected in all of the blow samples, and none were positive to Calnexin (Figure 3). Three out of four SEC derived-EV samples isolated from bottlenose dolphins’ blow and the one isolated from the long-finned pilot whale’s blow derived-EVs also resulted positive for OmpA, a membrane protein present in Gram negative bacteria and in the derived EVs (Fig 3). In contrast, none of the EV-samples isolated from bottlenose dolphin’s blow with UC showed any positive signal for the OmpA antigen (Fig 3).

**Figure 3.**
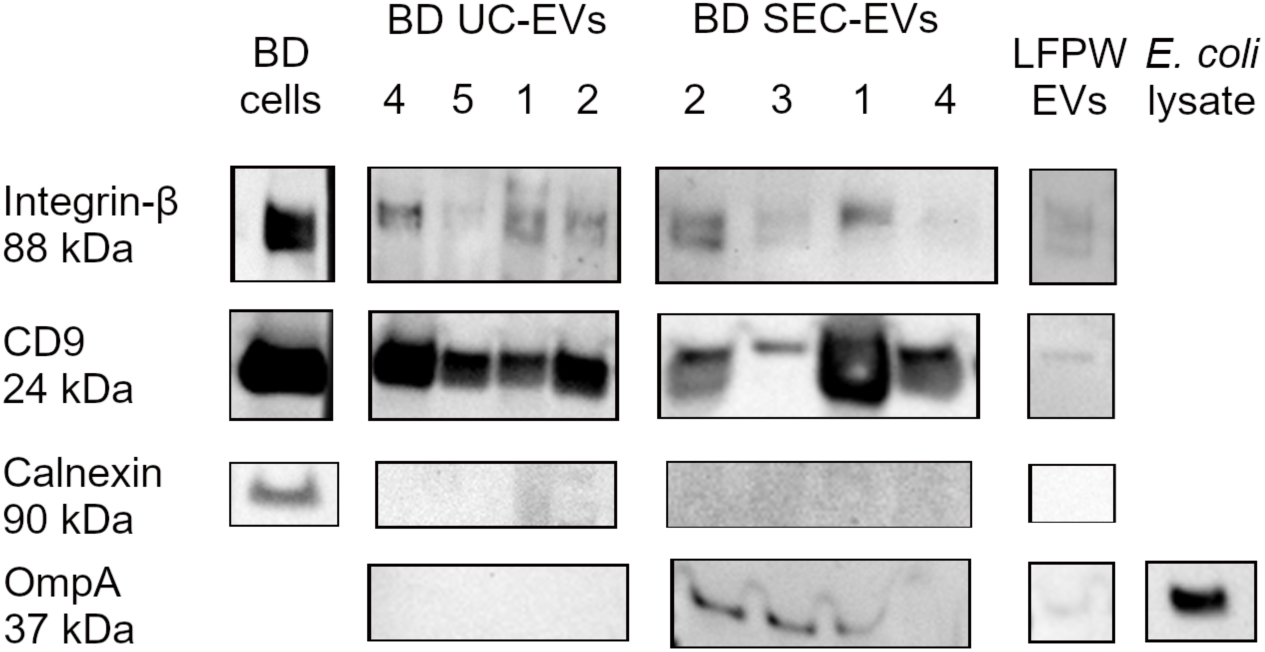
Western blot analysis of extracellular vesicles (EVs) isolated from bottlenose dolphin (BD) and long-finned pilot whale (LFPW) blow samples. Methods used for isolation were ultracentrifugation (UC) or size exclusion chromatography (SEC). The sample ID number is indicated on each lane. A fibroblast BD cell line (BD cells), and *E. coli* lysates are used as cellular positive controls. All EV-samples expressed the EV-markers integrin-β and CD9 and were all negative to the negative control Calnexin, a protein of the endoplasmic reticulum. Three BD SEC- (ID 2, 3, 1) and the LFPW-EVs were positive for the bacterial membrane protein OmpA.

## Discussion and conclusion

The potential diagnostic applications of biofluid-derived EVs have been largely explored for human medicine in the last two decades [29,43,45,52]. In recent years, this potential has shown up in veterinary medicine as well [47]. Nevertheless, EV-research in wild and exotic animal species is still largely unexplored, likely due to the practical and ethical challenges related to sample collection. In cetaceans, blow sampling represents a minimally invasive approach that allows the *in vivo* study of these animals. So far, however, the analysis of blow samples has been mostly limited to the assessment of hormones, bacteria composition or immune markers [16,17,20,25–27].

Here, we report for the first time the purification of extracellular vesicles (EVs) from cetacean blow samples. We conducted a methodological study in which two EV isolation techniques were applied and compared using blow samples collected from five captive-held bottlenose dolphins. The method demonstrating better results in protein analysis, was then selected to isolate EVs from a blow sample obtained from a wild long-finned pilot whale. This work serves as a proof-of-concept study to evaluate the feasibility of using blow-derived EVs to investigate the health conditions of wild cetaceans.

The two protocols tested on bottlenose dolphin blow samples were based on UC or on SEC[53]. EVs isolated by the two different methods gave different results in terms of particle concentration and protein analysis. At NTA, particle concentration resulted heterogeneous among EV-samples isolated by UC or SEC, varying from the order of magnitude of 10^8^ to 10^12^ particles/mL. One possible explanation for this heterogeneity, can be individual differences, such as sex and age, which are related to the different profile of EVs isolated from human biofluids [54]. A second, non-mutually exclusive, explanation is the difference in exhaled respiratory condensate volumes that were collected. Despite heterogeneity, the mean particle concentration was significantly higher in UC derived-EVs compared to SEC. This finding concurs with published studies that show UC tends to concentrate higher numbers of nanoparticles comparted to SEC [50]. Considering particle size, bottlenose dolphin UC and SEC-derived EVs were measured with NTA and AFM. While the size of EVs did not differ significantly between UC and SEC, NTA reported a mean diameter considerably larger (> 200 nm) compared to AFM (< 80 nm). This difference is probably due to some commonly reported limitations of NTA: i. NTA measurements are based on the Brownian motion principle, which can be influenced by the refractive index of nanoparticles, not allowing the analysis of all the nanoparticles present in heterogenous solutions at the same time; ii. The lower detection limit of the instrument does not allow to correctly detect particles < 50 nm; iii. The impossibility to distinguish EVs from non-EV particles or aggregates [55–57].

The canonical EV markers (CD9 and integrin-β) were detected in all of the EV samples. The presence of these proteins in bottlenose dolphin EVs had already been assessed in a previous study from our group, focused on the characterization of EVs released *in vitro* from a bottlenose dolphin fibroblast cell line [48]. In addition, all samples were negative to the negative control Calnexin, meaning the presence of EVs without a concurrent relevant cellular contamination in all samples [50]. Given that many studies have focused on analyzing blow samples for host-microbiome evaluations [17,23,58–60], we also investigated the presence of bacteria-derived EVs in cetacean blow samples to assess whether blow-derived EV analysis could be leveraged for these applications. OmpA, a membrane protein of Gram-negative bacteria and of the associated-EVs (called Outer Membrane Vesicles – OMVs) [61], was detected only in bottlenose dolphin SEC derived-EVs and not in UC derived-EVs. Although UC allows to collect a higher number of nanoparticles, SEC should allow to collect EVs in higher purity [50,53]. Therefore, the higher purity of EVs purified by SEC, together with the lower concurrent presence of non-EVs nanoparticles, might have allowed also the detection of Gram-negative bacteria or of their associated EVs among all the particles present in the EV-sample. The presence of Gram-negative bacteria in the blow-hole has been reported in several species of free-ranging and captive cetaceans and mainly included species of the phyla Proteobacteria and Bacteroidetes [23,58,62–65]. Of note, in Southern Resident Killer Whales and Humpback Whales, potentially pathogenic Gram-negative bacteria were described in blowhole samples (e.g., *Salmonella*, *Aeromonas*, *Fusobacterium*, *Pseudomonas*, *Shewanella* spp) [59,66].

To assess the feasibility of the isolation of blow derived-EVs in free-ranging animals during field conditions, we also characterized EVs isolated from a long-finned pilot whale blow sample collected with UAV. In this case, it was not possible to collect information on the age and sex of the specimen during the in-field sampling. SEC was selected as EV-purification protocol because of the limited volume of the blow sample and because, according to the literature, SEC should allow to collect EVs in higher purity [53]. At NTA, nanoparticles showed a mean diameter within EV-size range and a concentration sufficient for most EV-analysis but lower compared to the mean particle concentration of bottlenose dolphin blow derived-EVs purified with the same methodology. In addition, long-finned pilot whale EVs expressed the same EV-markers expressed by bottlenose dolphin SEC-derived EVs at WB. Among these markers, also OmpA was expressed, demonstrating the presence of bacteria derived-EVs also in this sample. In this case, the low particle concentration might be related to the different type of sampling: the use of drones and in field conditions probably allow the collection of less blow compared to manual sampling in animals kept under human care. Indeed, other studies have already described the low yield of biological molecules, particularly of RNA, in the blow sampled with drones from other species (*Megaptera novaeangliae)* and, in some cases, the need of pooling blow samples to gather reliable data [22,67]. However, another study performed on bottlenose dolphins described a sufficient amount of biological matrix in each single blow sample for microbiota analysis, suggesting that collection methods and in field conditions can cause great variations in the quantity of collected biological material [17].

In animals and humans, the microbiome of the airways is known to have a great influence both on health and disease [68]. The presence of bacteria derived-EVs in cetacean’s blow samples might allow, in future studies, to gain more information on their interaction with the host, which has not been fully characterized in these species yet. Although to date, it is not yet possible to fully discriminate between EVs released from the host or from bacteria, the analysis of EV-cargo, particularly of EV-related RNA, using in depth sequencing analysis may help identify their biological origin [69].

In humans, the analysis of the microbiome of the respiratory tract has been recently regarded as a good tool to detect biomarkers of inflammation [70]. The analysis of cetacean blow can be compared to the analysis of the bronchoalveolar lavage fluid (BALF) in terrestrial animals. The composition of BALF microbiome has been investigated in healthy dogs and cats, in dogs with bacterial pneumonia, in cats with chronic bronchitis or asthma, and in dogs after treatment with antimicrobial drugs, suggesting that diseases and the use of antimicrobial drugs affect the composition and diversity of lung microbiota [71–75]. In cetaceans, some studies analyzed the microbiome in blow samples but they mainly reported the composition in different individuals. Differences in microbiome complexity and composition have been detected in blow samples from Humpback whales before and after their migration period, supposing the correlation with fasting conditions [15,23,58,59].

Considering EVs derived from the respiratory tract, in animals they have been isolated and characterized only in horses, where EVs derived from the BALF have been demonstrated to have a distinct fatty acid fingerprint in horses with asthma compared to healthy ones [76]. In humans, the profile of BALF derived EVs has been explored instead also in other conditions [77]. For example, the miRNA profile of BALF derived-EVs in patients with acute distress syndrome, pneumonia and pulmonary sarcoidosis, resulted significantly different compared to healthy patients [78–80]. BALF derived EVs have also been described for their protein signature, which is different in patients with chronic obstructive pulmonary disease and, for their different lipid profile in cases of asthma [81–83]. However, to our knowledge, no study has investigated BALF derived-EVs in relation to host microbiome yet.

Considering the diagnostic potential of EVs isolated from human respiratory biofluids, blow derived EVs can represent a novel tool for investigating cetacean health conditions. In this preliminary methodological study, we demonstrate the feasibility of EV isolation from cetacean blow samples collected from both captive-held and wild cetaceans. The comparison of SEC and UC applied to bottlenose dolphin samples indicates that, although SEC yields fewer particles, it appears more effective than UC for the evaluation of protein marker presence, probably due to the higher purity of the EV sample. Therefore, we recommend carefully selecting the EV purification protocol in accordance with the final experimental objective. If a highly pure EV sample is required, SEC should be preferred over UC; conversely, UC may be more suitable when maximizing EV yield is the priority. To comprehensively assess the quality of EVs obtained with these methods, further studies including larger cohorts are needed and the analysis of EV associated molecules through RNA sequencing and proteomic analyses should be performed to perform in depth EV characterization. The quality of the final EV sample may also be influenced by the physicochemical properties of blow, such as osmolarity and pH. These factors were not assessed in the present study and may differ from those of other mammalian-derived biofluids [84], potentially requiring adjustments to EV purification protocols. Although the EV yield obtained here with both UC or SEC was sufficient for most characterization analysis, future work should also determine whether it is adequate to generate robust health-related information. Given the challenges of sampling wild cetaceans, investigations on animals under human care – where EV data can be correlated to known health parameters – will be valuable to provide insights on how blow derived EV composition varies with physiological or pathological states. Subsequently, samples from wild cetaceans should be integrated with contextual data, such as photo-identification catalogues and environmental data, to maximize individual-level information.

Developing minimally invasive, field-applicable approaches to assess the health status of live cetaceans represents a crucial advancement for marine mammal and ecotoxicology research. New approaches should overcome key limitations of *post-mortem* investigations, often hampered by tissue post-mortem lysis, and should enable *in vivo* monitoring of individuals and populations in their natural environment. Within this context, the characterization of blow-derived EVs emerges as a promising tool: by providing access to a complex and information-rich molecular cargo, EVs may reveal novel biomarkers of respiratory and systemic conditions, as well as responses to environmental stress. EV-analysis therefore has the potential to expand the diagnostic value of blow analysis, supporting more comprehensive health assessments in free-ranging cetaceans and contributing to a deeper understanding of the pressures affecting aquatic ecosystems.

## Acknowledgements

We thank the Italian Ministry of Environment and Energy Security and Costa Edutainment S.p.A. for the possibility of including the under human care bottlenose dolphins in the study. We also warmly thank Aurélie Célérier and Angelo Torrente for their support in organizing the field missions.

---

**S1 Video.** Video of the blow sampling from the long-finned pilot whale.

**S2 File.** Particle size in nanometers measured for each particle detected at Air-Atomic Force Microscopy for extracellular vesicles isolated with ultracentrifugation or size exclusion chromatography from bottlenose dolphin blow.

**S3 Figure.** Original uncropped figures of Western Blotting Analysis.

